# Genetic Correlates of Host Use in Scotland’s Pearl Mussels

**DOI:** 10.1101/2025.08.22.671716

**Authors:** Victoria L. Pritchard, Peter Cosgrove, Victoria Gillman, Kara Layton, Lydia McGill, Barbara Morrissey

## Abstract

The freshwater pearl mussel (*Margaritifera margaritifera*) is an ecologically important and highly endangered bivalve native to running freshwaters across Europe and eastern North America. Its life cycle includes an obligate parasite stage in which newly emerged larvae attach to the gills of juvenile salmonid fishes. In northern Europe, populations specialize on one of two hosts: Atlantic salmon (*Salmo salar*) or brown trout (*S. trutta*). Previous microsatellite studies of pearl mussels in the Nordic region have shown an association between host specialization and population genetic characteristics. Scotland is considered a remaining stronghold for freshwater pearl mussel, however current knowledge of genetic characteristics and host association of Scottish *M. margaritifera* populations is limited.

We combined minimally-invasive viscera swabbing with reduced-representation sequencing (nextRAD) to examine genetic diversity of pearl mussels at 5,486 genome-wide SNP markers across 18 populations in Scotland. Our results revealed a substantial variation among populations in genetic diversity and interpopulation differentiation which was strongly related to known host. Populations known to specialize on Atlantic salmon exhibited high genetic diversity (mean H_e_ = 0.24) and low inter-population differentiation (F_st_ = 0.026), even between rivers draining to opposite coasts. In contrast, populations known to specialize on brown trout or occurring where only trout are available consistently showed lower genetic diversity (mean H_e_ = 0.15 /0.14) and much higher inter-population differentiation (F_st_ = 0.160 /0.271), with many populations being highly genetically distinct even when geographically proximate. Principal component analysis and neighbor-joining trees confirmed this pattern, with salmon-specialist populations clustering together while trout-associated populations formed discrete, population-specific clusters.

These findings mirror previously observed patterns in other parts of the *M. margaritifera* range and indicate that population sizes and migratory behavior of hosts may drive contrasting evolutionary trajectories in pearl mussel populations. The striking genetic differences between salmon- and trout-specialist populations have important implications for conservation planning, as they indicate differential capacity for local adaptation and vulnerability to reduced population sizes. Our results suggest that population genetic characteristics could be used to predict host associations for unstudied populations, providing a valuable tool for conservation management. The study emphasizes the importance of considering both direct impacts on pearl mussel populations and the status of their salmonid hosts when developing conservation strategies for this rapidly declining species.

## Introduction

The freshwater pearl mussel (*Margaritifera margaritifera*, FPM) is a unioid bivalve mollusc that occurs in nutrient-poor, highly oxygenated, fast flowing freshwater across Europe and eastern North America. As a freshwater mussel, it is considered a keystone ‘ecosystem engineer’, improving water quality and transferring nutrients from the water column into the benthos via filter feeding, and modelling the substrate via bioturbation and shell formation (Vaughn 2018). Once widespread and occurring at high densities, FPM have dramatically declined over the past century as a result of human activity (Young *et al*. 2001). The species is now IUCN listed as ‘Critically Endangered’ in Europe. One characteristic of this decline is a lack of recruitment, meaning that even large and dense populations may entirely consist of mussels nearing the end of their 150+ year lifespan. The reasons for this are not fully understood (Österling et al. 2008)

In common with all unioid bivalves, FPM have a remarkable, partly parasitic, life cycle. Mussels take a decade or more to reach maturity and can then reproduce semi-annually for over a century (Bauer 1987). Female or facultatively hermaphroditic FPM are fertilized by water-borne sperm and subsequently release millions of larval offspring. These ‘glochidia’ are obligate parasites which encyst on the gills of nearby juvenile host fish and grow there for up to 11 months, increasing several-fold in size before excysting into the stream bed (reference). The parasitic stage is considered to be an adaption to gain nutrients and counteract downstream transport of sessile adults (Rock *et al*. 2025). Glochidial infection is associated with several negative effects on the host, including reduced fish growth rate and altered behaviour (Rock *et al*. 2025; Filipsson *et al*. 2016). In Europe the two known host fish of FPM are Atlantic salmon (*Salmo salar*) or brown trout (*S. trutta*) (Bauer 1987). Both of these host species have also declined over the past century, and insufficient numbers of larval hosts is one potential factor limiting pearl mussel recruitment.

There is evidence that different FPM populations are adapted to different fish host species, and that this host specialism has a genetic basis. Rather than active host choice, the adaptation appears to be mediated through the ability of glochidia to avoid the host immune reaction and encyst in the gill (Young & Williams 1984). In mainland northern Europe, *S. trutta* is used as the host where it is the only species available, but in the presence of both species *S. salar* is often the dominant host (e.g. Karlsson *et al*. 2014 but see Österling & Wengström 2015). Salonen *et al*. (2017) performed experimental work in which they exposed caged salmonids to glochidia from different Finnish pearl mussel populations. They demonstrated substantial differences between mussel populations in their ability to exploit salmon vs. trout hosts, which reflected the historical salmonid community composition in their native rivers. Wacker *et al*. (2019) undertook similar experiments using neighbouring pearl mussel populations which co-existed either with brown trout alone or with both hosts.

They found that larvae from the former populations almost exclusively infested brown trout, while larvae from the latter populations almost exclusively infested salmon. In Scotland, glochidia surveys have also indicated differential host fish specialization among pearl mussel populations (Hastie & Young 2001; Baum 2015, 2018, Clements *et al*., 2018). Nevertheless, populations may retain the ability to exploit either host; Marwaha *et al* (2021), in experimental work with a salmon-specialist population, showed that maternal parents produced offspring with varying propensities to infect both salmon and trout.

The isolation of both pearl mussel and salmonid host populations within river networks also raises the prospect of local adaptation, whereby glochida have evolved to encyst most successfully on local fish. This scenario would imply that genetic changes in the host population – for example due to fish stocking – could have a detrimental effect on local pearl mussel reproduction. However, there is a co-occurring selective pressure on the local fish population to resist glochidial infection, meaning that, depending on the evolutionary dynamics of the system, non-local fish could be better hosts. Adaptation of glochidia to local hosts has been tested experimentally by several authors (Taeubert *et al*. 2010; Österling & Larsen 2013; Taskinen & Salonen 2022) and no consistent adaptive pattern has been found; in some cases, glochidia perform better on a local host, but in others a non-local host appeared more suitable.

Studies using microsatellite genetic markers have repeatedly demonstrated a relationship between FPM host specialization, and population genetic characteristics, although this may partly be driven by differences in the size and accessibility of streams that can support trout versus salmon and thus the size of the potential host population. In Ireland and Norway, FPM populations specializing on salmon exhibit higher genetic diversity and lower inter-population differentiation than trout-associated populations (Karlsson *et al*. 2014, Geist *et al*. 2018). Results from Österling *et al*. (2020) suggest, however, that this may reflect host abundance and migratory characteristics rather than species *per se*: Swedish FPM populations specializing on either Atlantic salmon or anadromous brown trout (‘sea trout’), both of which perform a marine migration, show greater diversity and lower inter-population differentiation than those exploiting resident brown trout in smaller streams.

In the United Kingdom, FPM are listed as a Priority Conservation Species. Scotland is considered a remaining stronghold (Young *et al*. 2001) although the species is on a similar downward trajectory to the rest of Europe, and many extant populations are restricted to remote upland streams. U.K. conservation measures include assisted reproduction programmes - in which juvenile salmonids are artificially infected with glochidia, either at the natural FPM site or in dedicated captive breeding facilities – habitat protection and restoration, and measures to shield extant populations from illegal pearl fishing, a continuing cause of mortality. Only one study to date (Cauwelier *et al*. 2006) has examined the population genetic characteristics of FPM in the U.K. The authors applied nine microsatellite loci across 44 populations, including 27 from Scotland, found strong genetic differentiation among rivers, and inferred the presence of two phylogenetic groups. However, they did not explore associations with the glochidial host.

Here, we combine non-invasive genetic sampling with characterization of genome-wide genetic diversity to further investigate the population genetic characteristics of FPM in Scotland, with specific reference to host use. With continuing decline of Scottish pearl mussels, and a concurrent increase in conservation efforts, knowledge of what host pearl mussels are using and how this correlates with population genetic parameters is important to guide management actions.

## Methods

### Re-evaluation of salmonid host associations

Host associations of Scottish FPM populations are assessed by capturing co-occurring juvenile salmonids via electrofishing and counting glochidia on the gills. Hastie & Young (2001), Baum *et al*. (2015, 2018) and Clements *et al*. (2018) provide results of previous glochidia surveys. These original studies had widely varying sample sizes for the two alternative hosts at each site, used different measures and statistical approaches to infer host specialization, and in some cases assessed the same river at different time points. We therefore re-analysed the raw glochidia count datasets from these studies, using presence/absence of glochidia on gills as the indicative variable and applying a Chi-square test to examine deviations from the null expectation of a random distribution of infected individuals across the two species. For one additional site we used local knowledge on host specialization gained from conservation-focused artificial infections.

### Target sampling locations

FPM sampled for this study were a mixture of populations targeted for known host specificity and other populations sampled opportunistically during other monitoring efforts. Sample information, including vice-county location, is provided in Table 1 and Figure 1. Due to the risk to FPM from pearl fishing, exact sample locations are not provided. Two measures were used as broad indicators of population size at the sampling location. First, channel width at the collection site was measured from aerial photographs using an online tool (https://gridreferencefinder.com/). Second, we used counts from previous standardized population surveys provided in Watt *et al*. (2015) and interpretated by NatureScot to divide FPM populations into four abundance categories: 1. Very High (indicative: >500,000 estimated total mussels), 2. High, 3. Medium, and 4. Low (indicative: < 500 total mussels). We also inferred likely brown trout life history (resident vs. anadromous) at each sampling site from publicly available information on local fisheries management. The majority of FPM populations sampled were in physically separate river drainages, however some were in different tributaries or localities in the same river (Figure 1).

**Table 1:** Information on sampled populations. **Location:** Site code; **Sample type:** Visceral swab or mixed glochidia from a single female; **River width:** Channel width measured at sampling location from aerial photographs; **Initial n**: Number of individuals sampled; **Population size:** broad estimate of number of pearl mussels in the local population, obtained by multiplying mean transect counts in Watt *et al*. (2015), by approximate river area and assuming 10% FPM habitat suitability: Very large >>5000 mussels; Large > 5000, Medium ≈ 5000, Small <5000, Very small <<5000; **Post-QC n**: Number of samples that yielded genomic data; **Length:** Shell length of sampled individuals, mean and range; **Ref:** Reports re-analysed for host specialization: 1 Hastie & Young 2001; 2 Baum 2017; 3. Clements et al. 2018; **Infected salmon/total**: Total number of salmon with glochidia divided by total number of salmon examined, over all surveys; **Infected trout/total:** Total number of trout with glochidia divided by total number of trout examined, over all surveys; **χ2** Chi-squared statistic for deviation from expected numbers of infected salmon and trout assuming no host specialization; **p:** probability of observed value of Chi-squared, 1df; **Conclusion:** inferred host specialization, from χ2 results or other information.

| Location | Sample type | River width | Initial n | Population Size | Post QC n | Length (mm) - mean & range | Ref. | Infected salmon/ total | Infected trout/ total | $\chi^2$ | p | Conclusion |
| --- | --- | --- | --- | --- | --- | --- | --- | --- | --- | --- | --- | --- |
| EI1 | Swab | 30m | 15 | Very large | 3 | 109.4 (98-129) | 4 | na | na | na | na | Known salmon specialist from artificial infections |
| EI1 | Glochidia |  | 5 |  | 3 |  |  |  |  |  |  |  |
| ER1 | Swab | 18m | 9 | Small | 6 | 89 (75-104) | none | na | na | na | na | Host unknown; likely only trout available. |
| ES1 | Swab | 4m | 17 | Small | 6 | 91.3 (67-106) | none | na | na | na | na | Host unknown; salmon & trout available |
| NE1 | Swab | 10m | 10 | Large | 2 | 91.5 (82-111) | none | na | na | na | na | Host unknown; salmon & trout available |
| OH1 | Swab | 3m | 20 | Very small | 2 | 86.6 (71-102) | none | na | na | na | na | Host unknown; salmon & trout available |
| M1 | Swab | 2m | 10 | Very small | 0 | 89 (63-113) | none | na | na | na | na | Host unknown; salmon & trout available |
| M2 | Swab | 60m | 10 | Large | 0 | 119.5 (92-141) | none | na | na | na | na | Host unknown; salmon & trout available |
| M3 | Swab | 6m | 10 | Very small | 0 | 99.1 (69-123) | none | na | na | na | na | Host unknown; salmon & trout available |
| M4 | Swab | 8m | 4 | Very small | 0 | 125.5 (123-127) | none | na | na | na | na | Host unknown; salmon & trout available |
| M5 | Swab | 18m | 6 | Very small | 0 | 120.5 (115-127) | none | na | na | na | na | Host unknown; salmon & trout available |
| M6 | Swab | 25m | 11 | Large | 0 | 122.2 (106-139) | 1 | 53/53 | 12/12 | 0 | ns | No host specialism |
| WI1 | Swab | 1m | 15 | Small | 15 | 84.4 (69-97) | 2 | 0/0 | 25/39 | na | na | Only trout caught |
| WI2 | Swab | 4m | 15 | Small | 13 | 98.1 (84-121) | 2 | 0/0 | 49/129 | na | na | Only trout caught |
| WI3 | Swab | 9m | 15 | Large | 12 | No data | 1,2,3 | 164/285 | 1/42 | 22.1 | <0.001 | Salmon specialist |
| WI4 | Swab | 2m | 18 | Small | 9 | 86.5 (69-113) | 2 | 0/0 | 59/117 | na | na | Only trout caught |
| WR1 | Swab | 14m | 15 | Very Large | 10 | 90.3 (70-101) | 1,2,3 | 262/405 | 9/47 | 14.6 | <0.001 | Salmon specialist |
| WR2 | Swab | 9m | 15 | Large | 9 | 105.7 (56-130) | 2,3 | 3/116 | 11/54 | 14.2 | <0.001 | Trout specialist |
| WR3 | Swab | 7m | 15 | Large | 15 | 99.3 (85-126) | 3 | 0/122 | 4/21 | 23.2 | <0.001 | Trout specialist |
| WR4 | Swab | 4m | 15 | Medium | 11 | 88 (76-101) | 2,3 | 0/15 | 52/156 | 5 | <0.05 | Trout specialist |
| WS1 | Swab | 3m | 15 | Medium | 8 | 73.7 (59-89) | 2 | 0/0 | 22/80 | na | na | Only trout caught |
| WS2 | Swab | 2m | 15 | Small | 10 | 102.9 (86-115) | 2 | 0/0 | 36/50 | na | na | Only trout caught |
| WS3 | Swab | 24m | 15 | Medium | 8 | 76.6 (65-95) | none | na | na | na | na | Above waterfall: only trout available |
| WS4 | Swab | 4m | 16 | Large | 8 | 93 (68-109) | 2,3 | 0/281 | 34/50 | 191 | <0.001 | Trout specialist |
| WS5 | Swab | 27m | 14 | Large | 6 | 104.1 (83-124) | 2 | 47/131 | 0/3 | 1.1 | ns | Inconclusive; mostly salmon caught |
| WS6 | Swab | 35m | 10 | Large | 0 | No data | none | na | na | na | na | Host unknown; salmon & trout available |

**Figure 1.**
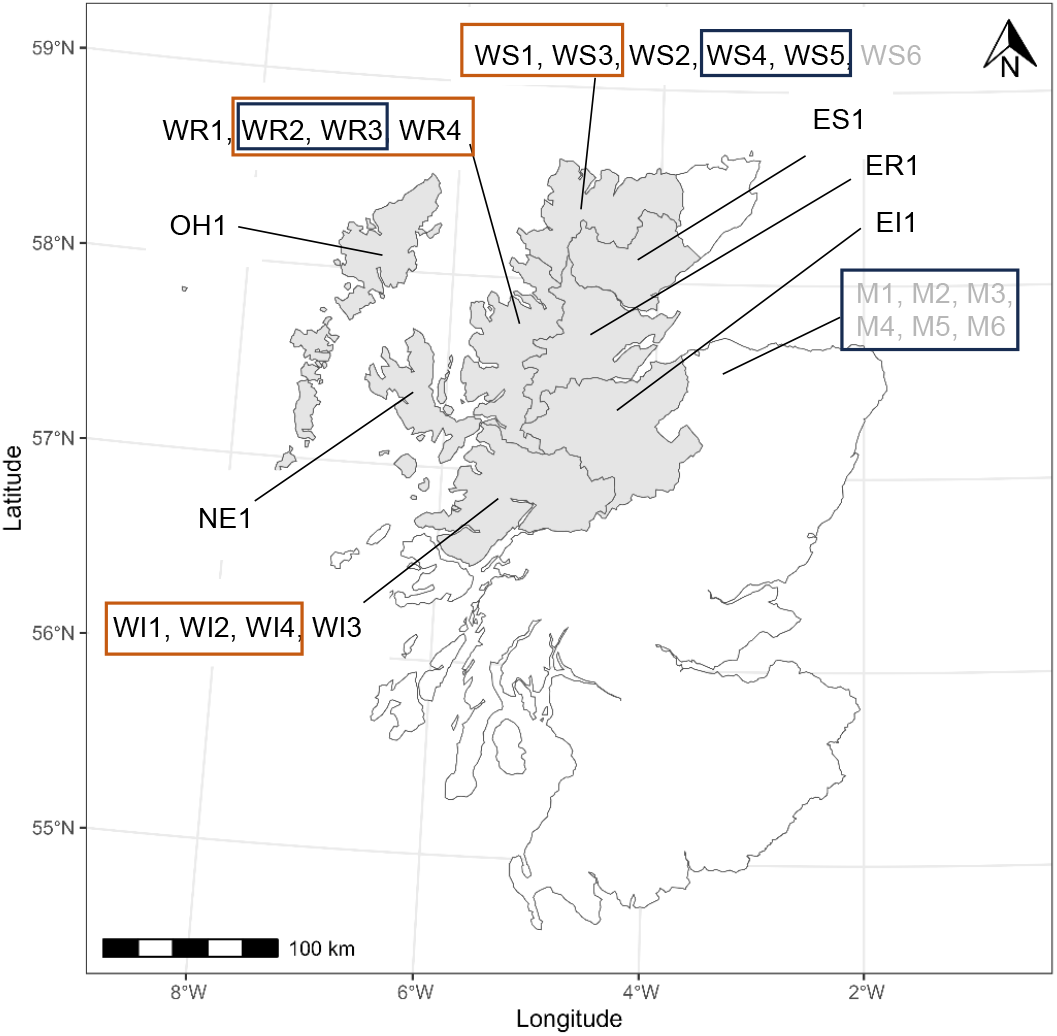
FPM sampling regions. Location IDs refer to vice-county of origin as follows: East Inverness (EI); Easter Ross (ER); East Sutherland (ES); North Ebudes (NE); Outer Hebrides (OH); West Inverness (WI); Wester Ross (WR); West Sutherland (WS). Black boxes indicate collection sites within the same river drainage. Orange boxes indicate collection sites in different drainages but within 5km of one another. Sites names in grey indicate that samples did not yield any useable genetic data.

### Sample collection

Samples for this study were collected during FPM survey or restoration work. DNA was obtained from live mussels in the field by viscera swabbing (Karlsson *et al*. 2013) using IsoHelix™ DNA buccal swabs. Briefly, the pearl mussel was temporarily removed from the water, the shell was slightly opened using modified pliers, and the swab was run across the visceral mass inside the mantle. Samplers recorded sample location and mussel shell length. Swabs were immediately immersed in 5ml BuccalFix™ buffer. Once returned to the laboratory, swabs and buffer were stored at 4°C until DNA extraction.

Additional samples of free-swimming glochidia, each produced by an individual female, were collected from one population during artificial infections; these were stored in 97% ethanol at 4°C.

### DNA extraction and quantification

DNA was extracted from swabs using Isohelix™ DNA Isolation kits, and from glochidia using Qiagen Blood & Tissue kits, both following manufacturers’ instructions. We assessed concentration and purity of DNA extracts via spectrophotometry using a QIAxpert microfluidic system (Qiagen). Any sample containing < 10ng/ul of DNA was excluded from further analysis. We assessed the presence of intact genomic DNA in the sample via gel electrophoresis: a 5ul aliquot of the extraction was combined with 1ul 6x loading dye and run along a 1% agarose gel at 50V for 2 hours, followed by visualization using SYBRsafe™ DNA stain. A clear band of high molecular weight DNA was considered indicative of a sufficiently non-degraded sample.

As gel visualization indicated substantial sample contamination with RNA and/or other non-target molecules, which frequently co-purify with DNA when dealing with mollusc tissue (Adema 2021), samples that passed initial quality control were subject to magnetic bead clean-up. Any sample with DNA concentration >40ng/ul was first normalized to 40ng/ul in a volume of 50ul by the addition of 10mM Tris-HCl using a dilution robot (QIAgility, Qiagen). Clean-up used Ampure XP beads according to the manufacturer’s protocol with a ratio of 1x sample to 0.4x beads (40ul:16ul), two rinse step each with 125ul 80% ethanol and a final elution with 20ul of 10mM Tris-HCl. QIAxpert and agarose gel QC steps were repeated for cleaned samples.

### NextRAD sequencing

DNA samples passing final QC steps were normalized to 150ng in 20ul volume of buffer and sent to SNPsaurus LLC, Eugene, Oregon for restriction site-associated DNA sequencing (nextRAD) following methods described in Russello *et al*. (2015). Briefly, genomic DNA was fragmented and adapter sequences ligated using Nextera DNA Flex reagent (Illumina, Inc), scaled for 50 ng of genomic DNA. DNA fragments containing the sequence GTGTAGAGCC were selectively amplified and the resulting nextRAD libraries were single-end sequenced on a Novaseq 6000 with one S1 lane of 122 bp reads (University of Oregon). SNPsaurus undertook two internal quality control steps. One sample was replicated during library generation to confirm genotyping repeatability. To check that the method was primarily amplifying pearl mussel DNA rather than non-target taxa, a random 1000 nextRAD sequences per individual were aligned against the full NCBI GenBank nucleotide database using BLAST (Altschul *et al*. 1990). Sequence reads were separated by individual-specific index tags and returned in fasta format.

### Variant calling and filtering

Untrimmed RAD sequence reads were aligned against the combined V2 *M. margaritifera* nuclear and mitochondrial genomes (GCA_029931535.1_MarmarV2 & NC_043836.1, Gomes-Dos-Santos *et al*. 2019; 2023) using bwa mem (Li 2013) with default parameters. Variant sites - single nucleotide polymorphisms (SNPs) and insertions/deletions (indels) - were called using bcftools mpileup followed by the multiallelic caller (Danacek *et al*. 2021), with read depth and genotype quality reported for each site. We did not downgrade mapping quality for reads containing excessive mismatches, as this can result in loss of data when the reference genome and experimental samples are from different populations (in this case Portugal vs. Scotland). Variants were called for all samples simultaneously.

We used vcftools (Danecek *et al*. 2011) to apply the following initial filters: minimum variant site quality (minQUAL) = 30, minimum depth of sequencing coverage at a variant site in an individual (minDP) = 20, min minimum genotype quality at a variant site in an individual (GQ)=20, maximum missing genotypes per variant site = 90%. We then imported the vcf file into PLINK 1.9 (Chang *et al*. 2015) and filtered further as follows: all indel and mitochondrial variants removed; maximum missing genotypes at a site = 20%; maximum missing genotypes per individual = 40%; minimum minor allele frequency = 0.02. We inspected the remaining SNPs for observed heterozygote excess across the entire dataset, indicative of technical genotyping problems, using the PLINK command --hardy. We corrected the significance threshold for multiple testing using the Benjamini & Hochberg (1995) procedure with a false discovery threshold of 10% and removed all SNPs that exhibited heterozygote excess and exceeded this threshold. Observed SNP heterozygote deficiency within a multi-population dataset can reflect either technical genotyping errors or population subdivision, and therefore we did not filter on this criterion.

Large numbers of closely linked markers can bias population genetic analyses. We therefore pruned SNPs that were in strong linkage disequilibrium (LD) with each other on the same genomic scaffold (r^2^ > 0.5) using the PLINK command –indep-pairwise 50 5 0.5.

### Statistical analysis

Using the LD-pruned dataset, we calculated average genome-wide observed and expected heterozygosity, inbreeding coefficient (F_is_), and mean proportion of missing genotypes for each sampled pearl mussel population from per-SNP values returned by PLINK (commands --family --hardy). To examine variation in levels of relatedness within populations, we used PLINK to separate each population, reapply the minimum minor allele frequency threshold, and estimate the kinship coefficient PI-HAT between each pair of individuals.

We used currentNe2 (Santiago et al. 2025) with the full SNP dataset to estimate contemporary effective population size (N_e_) at each sampling location. This approache infers N_e_ from observed LD between loci which are known to be weakly physically linked (Waples & Do 2009), and for our study we expect this estimate to be biased low because the fragmented FPM reference genome precludes identification of many physically linked marker pairs. Nevertheless, relative N_e_ estimates among FPM populations can still be informative. The few estimates of recombination rates in bivalves range widely from 0.7cM/Mb in *Mytilus* (Faure *et al*. 2008) to 3.05 cM/Mb in *Crassostrea* (Gagnaire *et al*. 2018) and we therefore applied the default rate of 1cM/Mb.

To explore genetic differentiation among sampling locations, we used PLINK (command – fst) with the LD-pruned dataset to estimate global F_st_, weighted for sample size (Weir & Cockerham 1984) across all sampling sites, and across sites with known salmon-specialist FPM, known trout-specialist FPM, and sites with only trout available. We used StAMPP in R (Pembleton *et al*. 2013) to estimate pairwise F_st_ among all locations.

We used three approaches to explore population genetic structure within the dataset without a-priori assumptions about its form. First, we used PLINK to perform a principal component analysis on the genome-wide identity-by state matrix generated from the LD-pruned dataset. We started with 20 principal components and reduced this by inspecting the scree plot of eigenvectors to identify the most suitable number. We plotted the position of individuals along each retained PCA axis using ggplot in R (Wickham 2016; R Core Team 2025). Second, we used the dist function of Adegenet in R (Jombart & Ahmed 2011) to generate a Euclidean distance matrix based on allele sharing among the individual genotypes, used the nj function to generate a neighbour-joining tree, and plotted the unrooted tree using ggtree in R (Xu *et al*. 2022).

## Results

### Sample collection

A total of 335 swab samples and 5 glochidia samples were collected from 25 sampling sites (Table 1, Figure 1). Broadly estimated from shell length, the age of the sampled mussels likely ranged from <10 years to many decades (Hastie et al. 2000). The smallest mussels were sampled at site WS1 and the largest at sites M1-M6. Most of the FPM populations sampled co-existed with both anadromous and resident host fish.

### Re-evaluation of host associations

Results of the re-analysis of combined data from Hastie & Young (2001), Baum *et al*. (2018) and Clements *et al*. (2018) for genetically sampled FPM locations are shown in Table 1. One or more glochidial surveys had been performed for 14 of the 25 sampling locations. For six of these locations, infection status of trout vs. salmon differed from null expectations, with four indicating trout-specialist FPM and two indicating salmon-specialist FPM. One did not show deviations from null expectations but fish catch was strongly skewed towards salmon, and one had 100% glochidia prevalence in both species. Only trout were caught at the remaining five locations. Of the non-surveyed locations, one is known from artificial infections to contain salmon-specialist FPM, and one is situated above a waterfall impassable to salmon. For this study, the host specialization at the remaining sites is considered ‘unknown’.

### DNA extraction, sequencing and quality control

Of the 335 samples collected, only 175 yielded DNA of sufficient quality and quantity to be sent for RAD sequencing. Samples failed QC due to insufficient initial DNA yield (7%), absence of clear genomic DNA band on initial gel examination (24.8%); and either or both following bead clean-up (17.2%).

Following sequence alignment, variant calling and QC steps, 21 out of the 175 genotyped samples exceeded the missing data threshold and were removed from the dataset. Most of these samples had been near the QC threshold for DNA, and all had a high number of BLAST hits against species other than pearl mussel in most commonly various species of the bacteria *Pseudomonas* and *Aeromonas*.

The final genetic dataset consisted of 153 swab samples and three glochidia samples, from 18 of the 25 sampling locations. These were genotyped at 5,486 single nucleotide polymorphisms distributed across 815 genomic scaffolds, with an overall per-site genotyping rate of 96.3%. A set of 3,456 SNPs was retained following LD pruning.

### Intra-location genetic diversity

Population genetic parameters were only estimated for the 16 sampling locations with sample sizes ≥6. For one of these locations (EI1), three of the six samples were mixed glochidia originating from a single female fertilized by multiple males, some of which may also have been swabbed.

Expected heterozygosity ranged from 0.08 to 0.25 (Table 2). Samples from the three known salmon specialist FPM locations exhibited substantially higher H_e_ (mean = 0.24) than the four known trout specialist populations (mean = 0.15) or the six populations coexisting only with trout (mean = 0.14). Observed heterozygosity was almost consistently lower than H_e_ resulting in positive estimated inbreeding coefficients for most populations (F_is_). Nevertheless, ten of the 16 populations showed a mean kinship estimate of zero. Discounting EI1, for which results may have been biased by the inclusion of both glochidia and swabs from their possible male parents (female parents were not genotyped), the highest mean kinship coefficients were observed in trout-using FPM populations in smaller streams.

**Table 2:** Population genetic characteristics of sampled populations. Location: Site code; Host: **Inferred host:** Host as inferred in Table1: salmon, trout, only trout available or unknown; **Host Life History:** Whether possible salmonid hosts are anadromous, resident or both; **Missing genotypes:** mean per-sample proportion of missing genotypes; **H**_**e**_, **H**_**o**_: Genome-wide observed and expected heterozygosity, calculated from 3,456 LD-pruned SNPs; **F**_**is**_: inbreeding coefficient, calculated as (H_e_-H_o_)/H_e_, **Mean PiHat:** mean genome-wide proportion of identity-by-descent, estimated as described in the methods; N_e_ : point estimate of effective population size, estimated from observed LD as described in the text; **N**_**e**_ **90% CI:** 90% confidence intervals of Ne estimates.

| Location | Host | Host Life History | # samples genotyped | Missing genotypes | H<sub>e</sub> | H<sub>o</sub> | F<sub>is</sub> | Mean PiHat | N<sub>e</sub> | N<sub>e</sub> 90% CI |
| --- | --- | --- | --- | --- | --- | --- | --- | --- | --- | --- |
| EI1 | Salmon | Anadromous (salmon) | 6 | 0.14 | 0.24 | 0.27 | -0.12 | 0.0255 | na | na |
| ER1 | Unknown | Likely resident (trout) | 6 | 0.02 | 0.15 | 0.12 | 0.19 | 0 | 21.5 | 11.9-38.8 |
| ES1 | Unknown | Resident & anadromous (salmon/ trout) | 6 | 0.02 | 0.19 | 0.17 | 0.10 | 0 | 35.7 | 17.7-72.2 |
| NE1 | Unknown | Resident & anadromous (salmon/ trout) | 2 | 0.08 | na | na | na | na | na | na |
| OH1 | Unknown | Resident & anadromous (salmon/trout) | 2 | 0.02 | na | na | na | na | na | na |
| WI1 | Only trout available | Resident & anadromous (trout) | 15 | 0.02 | 0.16 | 0.14 | 0.11 | 0.0107 | 48.5 | 33.7-69.7 |
| WI2 | Only trout available | Resident & anadromous (trout) | 13 | 0.03 | 0.16 | 0.14 | 0.16 | 0 | 102.9 | 58.9-180.1 |
| WI3 | Salmon | Anadromous (salmon) | 12 | 0.03 | 0.24 | 0.22 | 0.12 | 0 | 162.1 | 81.3-323.1 |
| WI4 | Only trout available | Resident & anadromous (trout) | 9 | 0.04 | 0.16 | 0.14 | 0.11 | 0 | 64.4 | 33.9-122.4 |
| WR1 | Salmon | Anadromous (salmon) | 10 | 0.03 | 0.25 | 0.21 | 0.15 | 0 | 163.6 | 52.9-202.7 |
| WR2 | Trout | Resident & anadromous (trout) | 9 | 0.04 | 0.18 | 0.16 | 0.10 | 0.0005 | 49.5 | 28.0-87.4 |
| WR3 | Trout | Resident & anadromous (trout) | 15 | 0.03 | 0.2 | 0.16 | 0.22 | 0 | 165.8 | 93.1-295.4 |
| WR4 | Trout | Unknown (trout) | 11 | 0.03 | 0.11 | 0.09 | 0.13 | 0 | 61.2 | 35.6-105.2 |
| WS1 | Only trout available | Likely resident (trout) | 8 | 0.05 | 0.08 | 0.08 | -0.02 | 0.0404 | 35.8 | 19.7-65.0 |
| WS2 | Only trout available | Mostly resident (trout) | 10 | 0.05 | 0.17 | 0.18 | 0.00 | 0.0081 | 134 | 62.2-288.8 |
| WS3 | Only trout available | Resident (trout) | 8 | 0.02 | 0.13 | 0.12 | 0.12 | 0 | 60 | 30.2-119.0 |
| WS4 | Trout | Resident & anadromous (trout) | 8 | 0.06 | 0.11 | 0.1 | 0.03 | 0.019 | 22.3 | 13.8-36.2 |
| WS5 | Unknown | Resident & anadromous (salmon/ trout) | 6 | 0.09 | 0.25 | 0.25 | -0.01 | 0 | 38.4 | 18.1-81.4 |

N_e_ was not estimated for E1 due to the mixed sample types. For the remaining 15 populations, point estimates of N_e_ returned by currentNe2 ranged from 21.3 (ER1) to 165 (WR3) (Table 2).

### Inter-location genetic differentiation

Weighted global F_st_ across the 16 sampling locations was 0.215. Across the known salmon specialist populations, F_st_ = 0.026; across the trout specialists F_st_ = 0.160; and across populations only co-existing with trout, F_st_ = 0.271. Pairwise F_st_ ranged from 0.010 (between WR2 and its tributary WR3) to 0.49 (between trout-associated populations WS1 and WS5, Table 3).

### Overall population genetic structure

All 18 populations were included in the examination of overall population genetic structure. Based on inspection of the scree plot, 10 principal components were retained (Figure 2). The first three axes accounted for the largest portion of total variation (16.9%). PC1 separated out a cluster of three populations in small streams < 3km apart where trout are the only available host (WI1, WI2, WI4). PC2 separated a cluster that included all known salmon-specialist populations, together with populations WS5 and NE1, whose host is unknown. With the exception of one river/tributary pair (WR2 & WR3), all other populations separated into discrete population-specific clusters along PC2 - PC10. The salmon-specialist/WS5/NE1 cluster was not further split along any PC.

**Figure 2:**
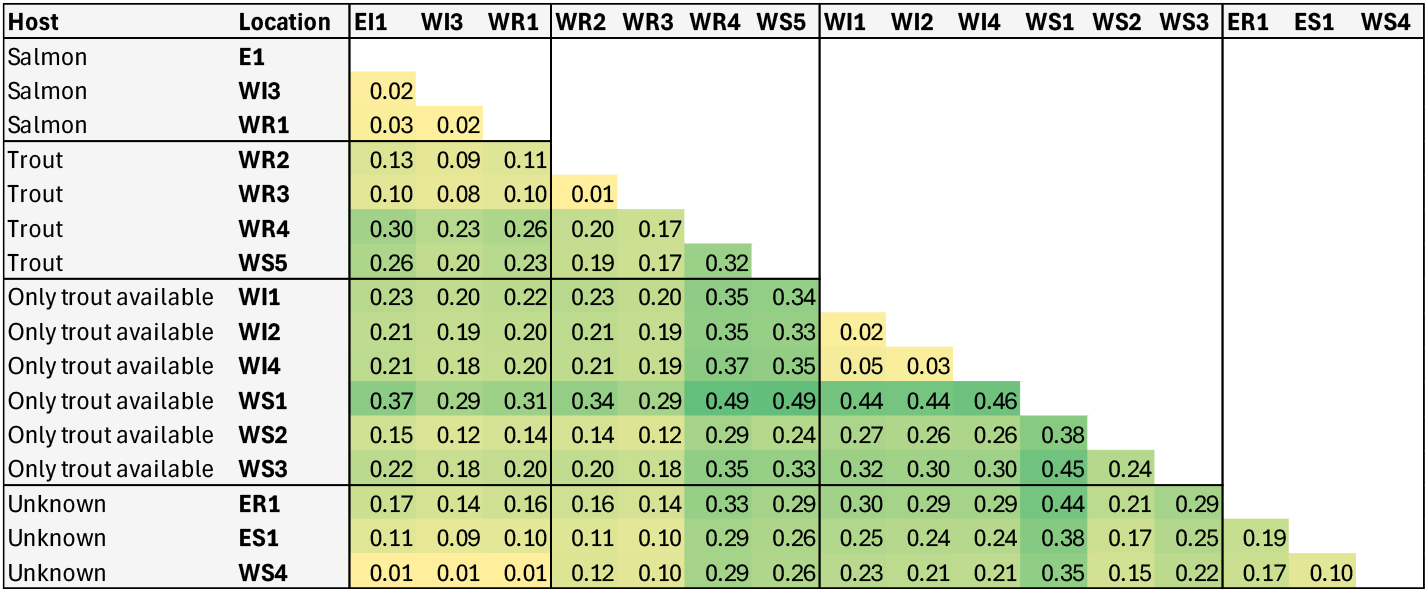
Pairwise F_st_ weighted for sample size (Weir & Cockerham 1984) between all locations with a sample size > 6. Locations are arranged by inferred glochidial host.

The neighbour-joining tree (Figure 3) revealed the same pattern; trout-only and trout-specialist populations broadly formed their own clusters, however individuals from known salmon-specialist populations plus WS5 and NE1, were interspersed within a single cluster.

**Figure 3:**
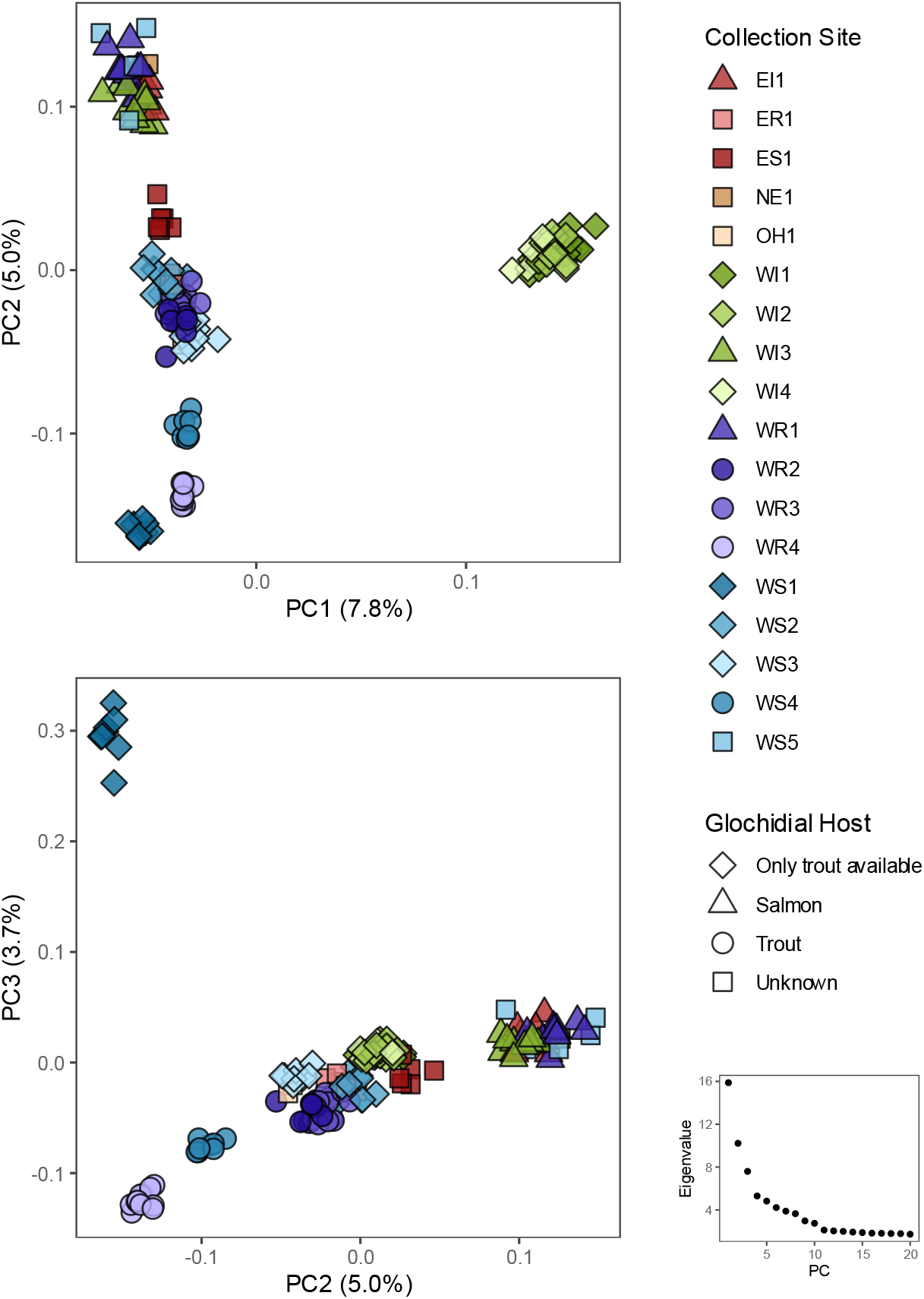
Results of Principal Component Analysis of the LD-pruned genomic dataset (3,456 SNPs, PC1-PC3). Each point represents an individual, plotted along the respective PC. Colour of point indicates sample location and shape indicates host specialization. Additional PCA plots (PC3-PC10) are provided in Figure S1.

**Figure 4:**
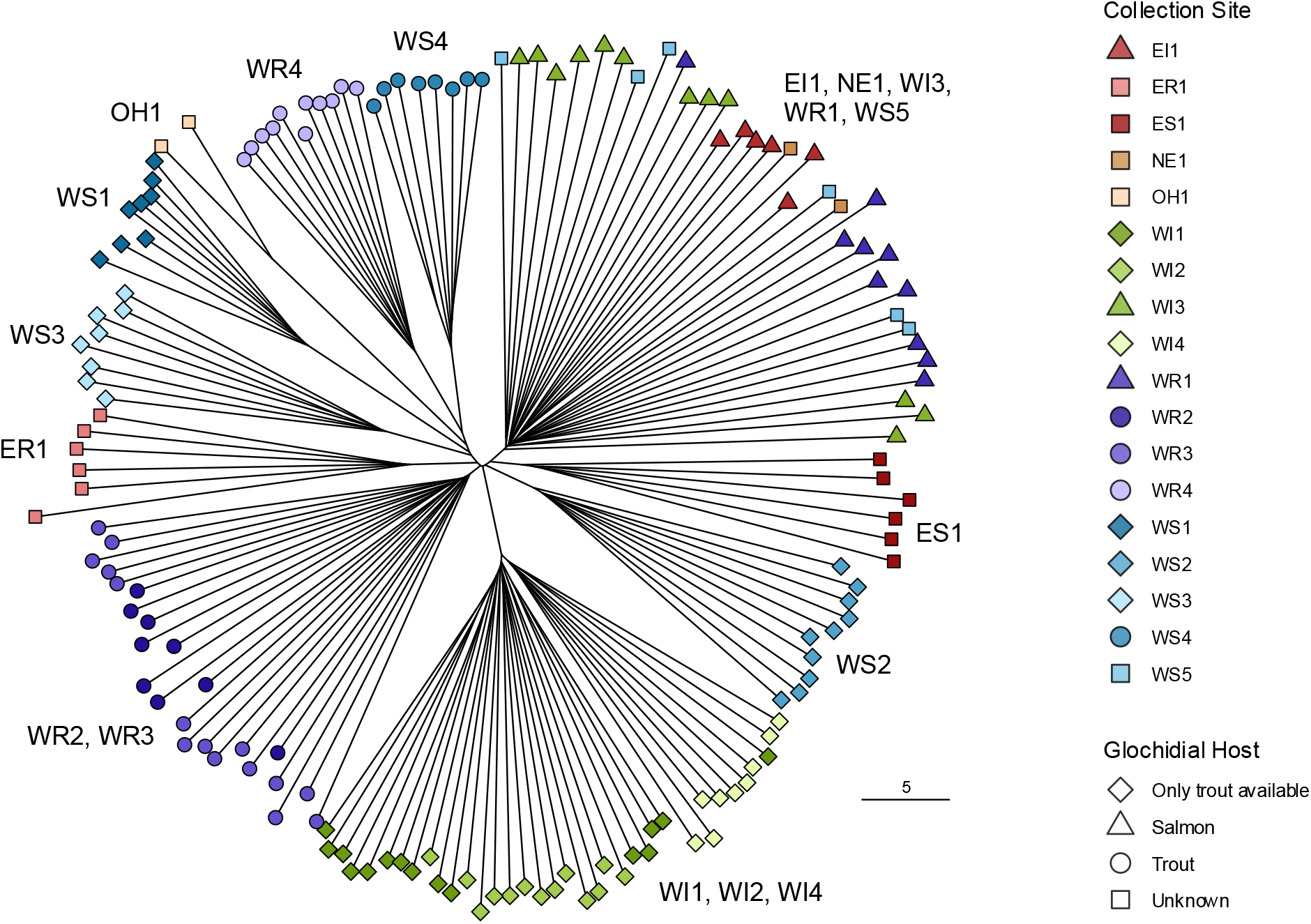
Unrooted Neighbour-joining tree based on allele-sharing distance among 156 individuals (3,456 SNPs). Point colour indicates collection site and shape indicates host specialization.

## Discussion

In this study, we combined non-invasive sampling with reduced-representation genome sequencing (nextRAD) to examine genetic diversity across FPM populations in Scotland. Our results revealed a strong association between salmonid host specialism, population genetic diversity and population differentiation. FPM specializing on Atlantic salmon exhibit relatively high genetic diversity and low genetic differentiation among populations. In contrast, FPM specializing on brown trout, or that occur in water bodies where Atlantic salmon are absent, have generally lower genetic diversity and much higher inter-population differentiation - so that many of these populations, including some geographically close to one another, are highly genetically distinct. Genotyped populations with unknown salmonid host associations exhibited population genetic characteristics typical of either salmon-specialist or trout-specialist populations, suggesting that these characteristics could be used to infer the host of the population without the need to undertake labour-intensive glochidial counts on fish gills.

Our findings closely mirror population genetic differences seen between salmon- and trout-specialist FPM in other parts of northern Europe (Karlsson *et al*. 2014, Geist *et al*. 2018, Österling *et al*. 2020). The very low differentiation among salmon specialist populations, even in rivers that drain to opposite coasts, is congruent with observations across the rest of the *M. margaritifera* range. For example, FPM show almost no genetic structuring across their entire distribution in North America, where Atlantic salmon may be the sole host (Zanatta *et al*. 2018). Other authors have attributed this pattern to glochidial transport among drainages by migratory hosts, which would presumably require glochidia to survive a marine migration. Glochida can remain encysted on Atlantic salmon gills through developmental adaptation to seawater (‘smoltification’) and for at least 50 days following transfer to the marine environment, however they are subsequently lost from the gills meaning that such transport could only occur in the earliest months of a salmon’s 1+ year marine migration (Treasurer & Turnbull 2000). Additionally, most trout-specialist FPM populations examined in this study co-occur with both anadromous and resident brown trout. This means that they could also have migratory hosts, although we do not have information about the relative proportions of the two trout life-histories at the various sites. An alternative explanation for lack of genetic differentiation among salmon-associated pearl mussel populations could be relatively recent post-glacial recolonization of rivers combined with very large effective population sizes and long generation times, meaning that allele frequencies have not yet diverged due to drift. However, although all known and putative salmon-specialist FPM populations in this study occur in relatively wider river channels and are recorded as having medium to very high abundance, there is no clear relationship between our (imperfect) LD-based estimate of N_e_ and known/putative host (Table 2). As Scottish freshwater pearls were historically an important trade commodity, we cannot completely rule out anthropogenic transfer of FPM as a contributor to the observed patterns – although historically this does not appear to have successfully established many new populations (Scherf 1980).

Our LD-based point estimates of N_e_ were in some cases orders of magnitude lower than population abundance estimates (e.g. the river from which sample WR1 was taken contains an estimated 0.6 to 1.2 million FPM (Watt *et al*. 2015) vs our N_e_ point estimate of 163.6). These estimates are strikingly similar to those obtained by Cauwelier *et al*. 2006 for some of the same FPM populations, using the LD approach with larger samples but only nine genetic markers. We strongly caution, however, that given the missing information on physical linkage among markers on different genomic scaffolds, the small and geographically limited samples used in our study, and the known low precision of LD-based N_e_ estimates for larger populations, especially when sample sizes are limited (Waples & Do 2009), that neither set of results should be taken as reliable indicators of true FPM N_e_/N_c_ ratios. We also observed an overall deficit of heterozygotes in at most sampling locations. Although this can result from uncorrected genotype calling bias, it is also a frequently observed feature of wild bivalve populations (Hollenbeck & Johnstone 2018). Rather than inbreeding, it is most commonly attributed to sweepstakes reproduction, where a few parents disproportionately contribute to recruitment success in each year, or selection occurring at early life stages. Nevertheless, estimates based on allele sharing (Table 2, PiHat) indicate relatively higher relatedness among FPM sampled from some trout-using populations in small streams. In particular, FPM from site WS1 show relatively high relatedness, low genetic diversity and particularly high differentiation from other populations. Even though the pearl mussel population at this site likely contains several thousand individuals, it has been heavily impacted by pearl fishing and contributed the smallest and thus youngest mussels sampled. Thus, these genetic characteristics may reflect ongoing recovery from a recent population bottleneck.

This work confirms that viscera swabbing is a suitable minimally invasive method of obtaining genomic DNA from pearl mussels in their native habitat (Karlsson et al. 2016). Extraction using a dedicated buccal swab kit followed by magnetic bead clean-up was able to produce DNA of sufficient quality for genome-wide genotyping using NextRAD. However, we were unable to generate data from 55% of the swab samples collected, most commonly because gel electrophoresis revealed no intact genomic DNA. Overall sample failure was not consistent across sampling sites, with some sites (M1-M6 and WS6) producing no useable samples and others (OH1, NE1) producing < 3 useable samples. Although swabs and DNA extractions had been stored for widely varying times before sequencing (7-26 months), in some cases exceeding manufacturer recommendations, we did not observe any relationship between storage duration and genotyping failure. We did, however, observe a correlation between sampler identity, sampling location and sample failure rate, suggesting that variation in swab handling and sampling methodology, or local water chemistry may contribute to failures. Blast hits of NextRAD sequences against microbial accessions suggest that visceral swabs could also be used to characterize aspects of the FPM microbiome, a potentially important aspect of bivalve health (Gignoux-Wolfsohn *et al*. 2024)

In summary, this first examination of genome-wide variation in Scottish freshwater pearl mussels has revealed a pattern of glochial host-associated population genetic structure, mirroring previous observations in northern mainland Europe. The striking contrast between population genetic characteristics of salmon- and trout-specialist FPM means that they are likely to have differential capacity for local adaptation to the host population and differential vulnerability to genetic risks associated with reduced population sizes, both of FPM themselves and of the host. Our results can both aid FPM conservation – by genetically predicting host associations for populations for which glochidial surveys have not been performed – and emphasize the importance of host populations in the conservation of FPM populations. The genetic composition and abundance of Scottish brown trout and Atlantic salmon populations have changed substantially over the past century due to human impacts that include habitat loss, fishing, climate change, deliberate stocking and accidental escapes of aquaculture fish. As previously emphasized by other authors (e.g. Hastie & Cosgrove 2001; Geist & Kuehn 2008), both direct impacts on FPM populations and interactions with their glochidial hosts should be considered when planning FPM conservation actions.

## Acknowledgements

This research was funded by NatureScot and UHI Inverness. We thank Iain Sime (NatureScot) for initiating the project and collecting samples, Chris Daphne, Corrina Mertens, and Orla Hilton for additional sample collection, Mark Coulson & Jenny O’Dell for contributions to project development, and Dasha Svobodova for laboratory help. Paul Etter and Eric Johnson at SNPsaurus LLC provided excellent nextRAD sequencing services.

## Author Contributions

Project development: VP, BM, PC

Sample collection: PC

Laboratory analysis: VP, BM, VG, LM

Bioinformatics: VP

Data analysis: VP Visualization: VP, VG

Manuscript drafting: VP

Manuscript reviewing: All authors.

---

**Figure S1:**
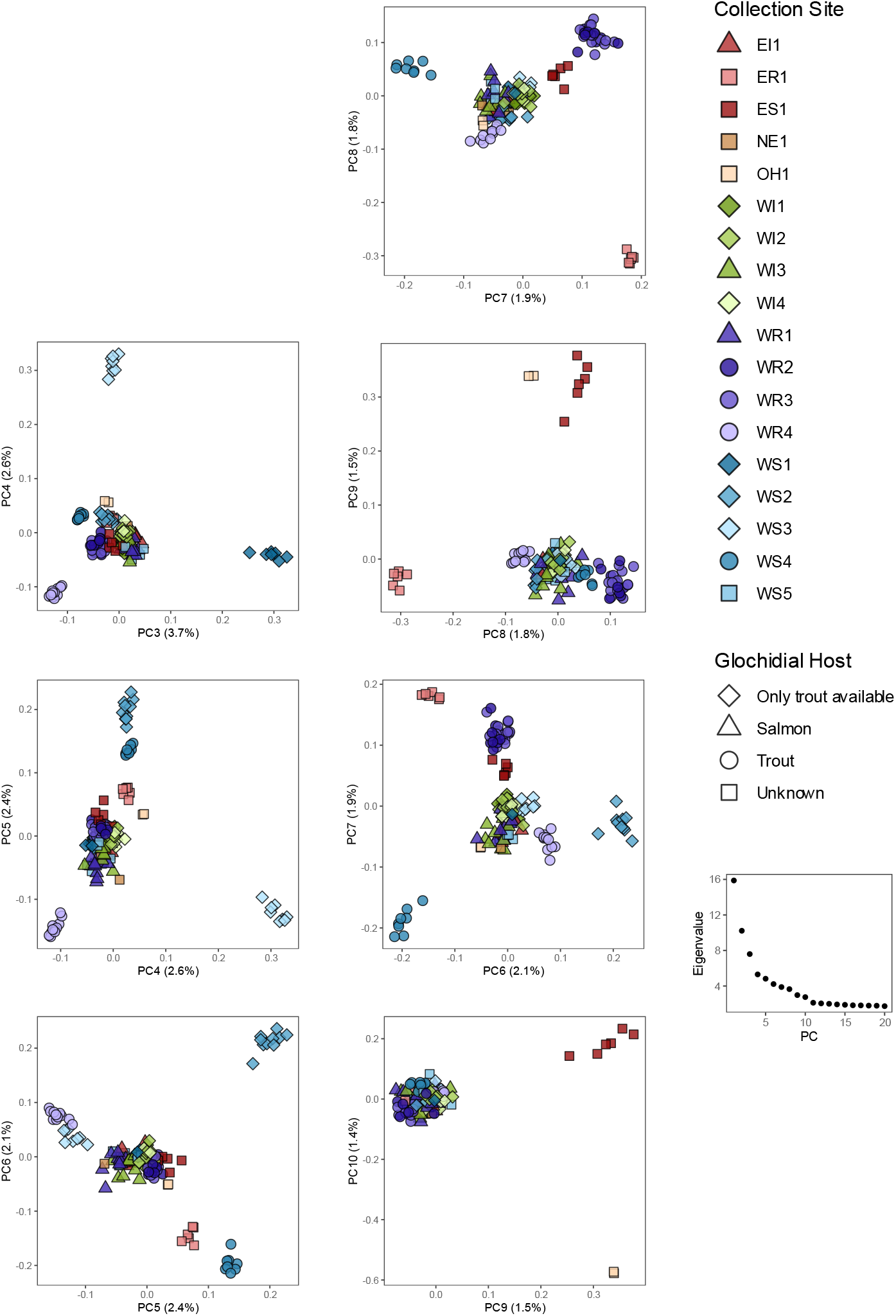
Results of Principal Component Analysis of the LD-pruned genomic dataset (3,456 SNPs, PC3-PC10). Each point represents an individual, plotted along the respective PC Colour of point indicates sample location and shape indicates host specialization.

